# Variations in neonatal age segments mortality in Kenya’s malaria epidemiological zones by community uptake of iron-supplements and anti-malaria drugs during pregnancy: Analysis based on 2014 Kenya demographic and health survey

**DOI:** 10.1101/388652

**Authors:** Boniface O. K’Oyugi

## Abstract

**Introduction:** Although past studies have established that iron-supplements and anti-malaria drugs taken by mothers during pregnancy reduce the risk of neonatal deaths in high prone malaria areas, little is known about their impact on mortality risks in neonatal age segments in Kenya. The study objective was to analyse variations in neonatal age segments mortality rates by uptake of these two antenatal care services and determine their effects on the age segments mortality in Kenya’s malaria zones.

**Data and methods:** This study used data from the 2014 Kenya Demographic and Health Survey (KDHS). Survival status information for 20,794 children born less than 60 months prior to interview date and reported mothers’ uptake of iron-supplements and anti-malaria drugs during last pregnancy was analysed. Life table method was used to estimate mortality rates and Poisson multivariate regression models were fitted to determine relative risks of death for the study variables.

**Results:** The results show that variations in neonatal age segments mortality in Kenya’s malaria zones are statistically insignificant. The contributions of early neonatal (0 to 7 days) to neonatal mortality rate are 80% and 100% in low and high malaria zones, respectively. Combined high community uptake of iron-supplements and anti-malaria drugs during pregnancy reduce significantly mortality risk in late neonatal (8 days to less than one month) in all malaria zones when effects of other risk factors are controlled for.

**Conclusions:** The findings suggest that future decline in neonatal mortality in all Kenya’s malaria zones depend mainly on reduction of early neonatal mortality. High community uptake of iron-supplements and anti-malaria drugs during pregnancy has significant reduction effect on late neonatal mortality in all malaria zones. This study recommends improvement of future KDHS data quality, especially on care for small and sick neonates.

## Introduction

Neonatal death is defined, according to the World Health Organization (WHO), as death within 28 days of birth of any live-born baby regardless of weight or gestation age [1]. Globally, about 2.6 million children died in the first month of life in 2016 with 1 million (38%) dying in the first day and another 1 million (38%) dying within the next six days [2]. Although the global neonatal mortality rate fell from 37 deaths per 1,000 live births in 1990 to 19 in 2016, this 49% decline was slower than the 62% mortality decline among 1-59 months old children [3]. In comparison to other regions of the world during the same period, sub-Saharan Africa had the slowest neonatal mortality decline of 40% but with impressive mortality decline of 63% among the 1-59 months old children [3]. Kenya’s neonatal mortality declined from 33 per live births 1,000 in 2003 to 22 in 2017 with 71% dying in the first week of life [4]. In similarity with sub-Saharan Africa region’s pattern, Kenya’s 33% reduction of neonatal mortality was slower than 63% mortality reduction among the 1-59 months old children during the period 2003 to 2014 [4–5].

About three quarters of the global neonatal deaths in 2016 were attributed to causes that are readily preventable and treatable with cost-effective interventions covering the antenatal period, time around birth and first week of life and care for small and sick neonates [3,6–8]. Attainment of their universal coverage (99%) would avert 70% of neonatal deaths with antenatal period interventions contributing about one third [3,6–8]. In 2016, the three main causes of neonatal deaths with contributions estimated at 35%, 24% and 15% respectively were: preterm birth complications (prematurity and low birth weight); intrapartum related conditions (birth asphyxia and birth trauma); and, sepsis (neonatal infections) [3,6–8]. In early neonatal period (0-6 days), preterm birth complications and intra-partum related conditions were the main causes of death (40% and 29% respectively) while sepsis and preterm were most dominant (44% and 21% respectively) in the later neonatal period (7-28 days) [3,7–8]. Sub-Saharan Africa region has the highest preterm birth complications [3,7–8].

Although antenatal period interventions have the least impact on neonatal deaths compared to the other two broad categories of interventions, they provide opportunity for integrated service delivery for pregnant women [9–12]. Antenatal period interventions enhance prevention, detection, and treatment of conditions or diseases associated with high risk of neonatal death during pregnancy, delivery, and care of new-born [9–12]. In developing countries, antenatal period interventions include provision of iron-supplements during pregnancy for early neonatal mortality reduction by reducing risk of preterm and birth asphyxia [10–15]. Iron deficiency is the most common cause of anaemia in pregnancy and iron-supplements are recommended for prevention [10,12]. About 20% of early neonatal deaths in Indonesia could be attributed to lack of iron-supplements during pregnancy [14,16].

Kenya’s 2014 uptake of iron-supplements during pregnancy was 69% with wide regional variations ranging from 40% in North Eastern province to 83% in Nyanza province [4].

Appropriate intermittent preventive treatment of malaria during pregnancy is also one of the antenatal period interventions associated with neonatal mortality reduction especially in high malaria endemic regions [14,16]. Malaria is a risk factor for maternal anaemia during pregnancy which is associated with increased risk of low birth weight and preterm delivery [17–19]. Intermittent preventive treatment of malaria in pregnancy reduces antenatal parasitaemia and placental malaria and its uptake is associated with 29% reduction in low birth and 31% reduction in neonatal mortality [10]. The adverse effects of malaria on risk of neonatal death are amplified when pregnant women are iron deficient [19]. Malaria transmission rate in new-born is estimated to be up to 20% in sub-Saharan Africa region [20]. Combined iron-supplements and malaria prophylaxis during pregnancy reduced significantly neonatal mortality in 19 malaria-endemic countries in sub-Saharan Africa [14,16].

More than 70% of Kenya’s population is at risk of malaria morbidity with pregnant women and children under age 5 years being the most vulnerable to infection [21]. Kenya has four malaria epidemiological zones: endemic covering the Lake Victoria in western Kenya and coastal region; highland covering western highlands of Kenya; semi and seasonal covering northern and south-eastern regions of Kenya; and, low risk covering central highlands of Kenya and Nairobi [22]. Neonatal mortality rates in Kenya vary by administrative region but do not depict clear patterns based on malaria epidemiological zones [3]. Uptake of intermittent preventive treatment of malaria in Kenya’s malaria endemic zones during pregnancy is 38% (based on WHO recommended standard of three or more doses) with wide variations by malaria transmission intensity ranging from 43% in coast endemic to 15% in highland epidemic [21–22].

Since most of the neonatal deaths occurring globally and in sub-Saharan Africa countries are attributable to causes which are readily preventable and treatable with known health care interventions, studies on impact of neonatal period interventions on neonatal deaths adjusted for other known mortality risk factors during: time around birth and first week of life; and, care especially for small and sick neonates [13–16,23–24]. These studies used mainly the Mosley and Chen’s 1984 conceptual framework for determinants of child survival in developing countries to identify mortality risk factors whose effects needed to be adjusted for [24–30]. The Mosley and Chen’s framework groups mortality risk factors into two broad groups: socio-demographic and birth characteristics; and, antenatal, delivery and post-delivery care services [31].

Among the numerous socio-demographic neonatal mortality risk factors commonly used as controls in studies undertaken in sub-Saharan Africa and south Asia regions, less than ten had significant effects and they included: first born infants; short birth intervals of less than 2 years; maternal age at delivery of the child of less than 20 or 30 plus years; male infants; perceived smaller than average sized infants; multiple births; high HIV/AIDS prevalence; and, rural residence [14,16,24,29–30]. The antenatal, delivery and post-delivery neonatal mortality risk factors found to have significant effects in south Asia and sub-Saharan Africa regions included: caesarean mode of delivery; and, delayed breastfeeding initiation period of beyond 24 hours [26–30]. Some of the neonatal mortality risk factors used as controls and found not to have significant effects, in this second broad group, included: non facility delivery; non-skilled delivery attendance; and, inappropriate immunization and treatment of sick infants [26–30].

Estimates of neonatal mortality rates and effects of iron-supplements and intermittent prevention and treatment of malaria in pregnancy on neonatal mortality have been reported in recent studies undertaken in south Asia and sub-Saharan Africa regions [14,16,24,32–35]. However, most of these studies treated neonatal period as a single age block and ignored the non-uniform shape of age pattern of mortality during neonatal period. In Kenya, provision of iron-supplements and anti-malaria drugs during pregnancy are part of the Antenatal Care (ANC) services package. The main objective of this study was to analyse variations in neonatal age segments mortality rates in Kenya’s high and low prone malaria areas by uptake of these two ANC services at community level. The secondary objective was to determine the effects of community uptake of these ANC services on neonatal age segments mortality risks in Kenya’s high and low prone malaria areas. Deeper understanding of neonatal age segments mortality risk reductions associated with uptake of these ANC services in Kenya would contribute to improvements in neonatal survival programmes.

## Data and methods

### Data

This study used data from the 2014 KDHS which was made available for public use including research during the official launch of its final report in December 2015 by the Kenya National Bureau of Statistics (KNBS). The 2014 KDHS dataset was also distributed to key stakeholder institutions and special interest groups. It can also be accessed, with official request, from KNBS and The DHS Program, ICF International. The 2014 KDHS is a nationally representative household survey data collected using the fifth National Sample Survey and Evaluation (NASSEP V) master frame which is operated and maintained by KNBS. The 2014 KDHS dataset consists of five separate recode data files namely: household; woman; male; couple; and, child. This study used the child data file which contains information on individual children born to women during the period 0 to 59 months before the interview. Appended to individual child’s record is information associated with the child gathered using household, woman’s and man’s questionnaires. This analysis was based on a total of 20,794 children born less than 60 months prior to interview date and these excluded interview month births.

### Ethical statement and data availability

The 2014 KDHS was approved by the Scientific and Ethical Review Committee of Kenya Medical Research Institute (KEMRI). Individual survey respondents agreed to participate voluntarily. Consent note was read to all respondents and signed by interviewers. The 2014 KDHS data released to researchers by KNBS and used in this analysis is without access to any personal data. The data can be accessed, upon official request, from KNBS (email: website: www.knbs.or.ke) and The DHS Program, ICF International (email: website: www.DHSprogram.com).

### Definitions and measurements

#### Outcome event

The outcome event in this analysis was the occurrence of death during neonatal period which was segmented into three age segments to capture the age pattern of neonatal mortality. The three age segments were: less than 1 day; 1 to 7 days; and, 8 days to less than one month. Two variables were used to measure neonatal age segment mortality. Life table probability of dying from birth to end of age segment per 1,000 live births was used to measure mortality rate. This measure facilitated analysis of mortality variations in malaria zones in Kenya by the study ANC services. Risk of death in age segment was used as dependent variable in the multivariate regression models fitted to determine effects of the study variables on neonatal age segment mortality.

Computation of life table probability and risk of death in neonatal age segment requires information on deaths and exposure days. The following information in the child data file were extracted: age at death in days for a neonatal death; age at death in completed months for a child who died after neonatal period; and, age in completed months for a child who was alive on interview date. A child who was alive on interview date, with age in completed months recorded as zero, was assumed to have lived for 15 days. A child reported to have died on the day of birth was assumed to have contributed a half exposure day in the first age segment. Total number of deaths in a neonatal age segment was obtained by summing all deaths in the age segment. Total exposure days in a neonatal age segment was the sum of total exposure days contributed by children who died in the age segment and total exposure days contributed by children who lived beyond the age segment. A child who lived beyond the age segment contributed full exposure days in the age segment.

#### Study antenatal care services

The ANC services of interest to this study were uptake of iron-supplements and anti-malarial drugs during pregnancy. Proportions of children in 2014 KDHS sample clusters whose mothers took each of these two ANC services were applied in this analysis as their community uptake estimates. Unfortunately, antenatal care information for woman’s uptake of iron-supplements and anti-malaria drugs was only captured for last pregnancy in 2014 KDHS. This resulted into a large proportion of children in this analysis with missing information on their maternal uptake of these services during pregnancy (66% for iron-supplements and 29% for anti-malarial drugs). Children with missing information were mainly cases of non-last-born children and also last-born children who had died prior to interview date. Weighted proportions of survey sample clusters’ uptake values for each study ANC service were computed to resolve the problem of sample variability at cluster level. The distribution of weighted clusters’ uptake proportions for each study ANC service was arranged in ascending order and a 40:60 percentile criterion used to group obtained weighted proportion into low (if 40th percentile or less) and high (if 60th percentile or more).

#### Malaria epidemiological zones

The 2014 KDHS sample clusters identification numbers were used to categorise them into malaria zones. Both the 2014 KDHS and 2015 Kenya Malaria Indicator Survey (KMIS) were conducted using the NASSEP V master sampling frame. Appendix A of the 2015 KMIS Report contains distribution of malaria epidemiological zones and sample allocation clusters by the 47 counties in Kenya [22]. In this analysis, the four Kenya’s malaria epidemiological zones were further grouped into high prone malaria areas (comprising of endemic and highland) and low prone malaria areas (comprising of semi, seasonal and low risk).

#### Confounding neonatal mortality risk factors

This study used the 1984 Mosley and Chen’s conceptual framework for analysis of determinants of child survival in developing countries to identify potential neonatal period mortality risk factors that needed to be controlled for. This framework classifies the risk factors into two groups: socio-demographic and birth characteristics; and, antenatal, delivery and post-delivery care services. During the preliminary stages in this analysis, only significant neonatal mortality risk factors derived from literature review with a bias on recent studies in south Asia and sub-Saharan Africa regions were considered for inclusion. In the final stages in this analysis, only those found to be significant at 95% confidence level and above, in bivariate regression analysis for neonatal age segment mortality determinants, were included as control variables. Brief descriptions and categorisations of control variables used in each neonatal age segment mortality analysis are provided below.

In age segment period day of birth, the following three socio-demographic and birth characteristics factors were used as confounding mortality risk variables: HIV positive pregnant women per 10,000 population (low if < 14.4 and high if ≥ 14.4); maternal age at child’s birth in years (low risk if 18-34 and high risk if < 18 and 35+); and, birth type (low risk if single and high risk if multiple). Only one mortality risk factor associated with antenatal, delivery and post-delivery care services was used as control variable and this was proportion in community with early breastfeeding initiation period of less than 24 hours (low if < 62.5% and high if ≥ 62.5%).

In age segment period 1 to 7 days, five socio-demographic and birth characteristics factors were used as confounding mortality risk variables namely: HIV positive pregnant women per 10,000 population (categories same as in the first age segment); preceding birth interval length in months (short if < 12 and long if first birth or 12+); birth type (categories same as in first age segment); elevated birth order (high if birth order =1 or 4+ and maternal age at child’s birth < 18 or 35+ years; and, low if birth order is 2-3 and maternal age at child’s birth is 18-34 years); and, maternal education (high risk if none or primary; and low risk if sec+). In this age segment two mortality risk factors associated with antenatal, delivery and post-delivery care services were used as control variables. These were: proportion in community with required tetanus injections during pregnancy (low if < 55.6% and high if ≥ 55.6%); and, mode of delivery (not caesarean section and caesarean section).

In age segment period 8 days to less than 1 month, only one socio-demographic and birth characteristics factor was used as control variable and this was HIV positive pregnant women per 10,000 population (categories same as in the first two age segments). Two mortality risk factors associated with antenatal, delivery and post-delivery care services were used as control variables in this age segment. These were: birth type (categories same as in first two age segments); and, proportion of children sick with diarrhoea in community who received appropriate treatment/care (low if < 55.6% and high if ≥ 55.6%).

The 2014 KDHS data file doesn’t contain information needed to compute survey clusters’ values of HIV positive pregnant women per 10,000 population. Administrative county level values were therefore used in this study and were computed using information contained in following two published reports available for public use: 2014 Kenya HIV County Profiles (KHCP); and, Kenya Population Projections (KPP) [36–37]. The 2014 KHCP Report contains numbers of HIV positive pregnant women in the year 2013 for each of the 47 administrative counties in Kenya. The 2010 KPP Report contains 2013 population projections for each of the 47 counties. Fortunately, the NASSEP V frame, which was used in the 2014 KDHS, had survey clusters covering all the 47 counties. The individual child information in the child data file was linked to the computed county level HIV positive pregnant women per 10,000 population using survey county codes. In this analysis, the computed county level estimate was used as the measure of this control variable.

## Methods

This study applied life table and multivariate Poisson regression methods. Life table method was used to calculate probabilities of dying from birth to age x in days per 1,000 live births (_0_*q_x_*,). The computed _0_*q_1_*, _0_*q_7_* and _0_*q_30_* are probabilities of dying by ages 1 day, 7 days and 1 month, respectively per 1,000 live births. These probabilities were computed for each malaria prone area by study ANC service. Detailed technical expositions of life table method are available in the book published by International Union for the Scientific Study of Population (IUSSP) [38]. The Statistical Package for Social Sciences (SPSS) was used to compute total deaths and total exposure days in neonatal period age segment in malaria prone area by study ANC service. 95% Confidence Intervals (CI) for the probability values were also computed using the package.

Multivariate Poisson regression models were fitted to estimate effects of the study ANC services on neonatal age segment mortality in each Kenya’s malaria prone area when controlling for effects of potential confounding mortality risk factors. In this analysis, the inclusion criterion was age segment mortality risk factor found to be significant at 95% confidence level and above in bivariate regression analysis. Poisson regression was chosen due to its ability to generate odds ratios estimates for covariates and also deal with any violation of the underlying assumption on equal mean and variance in distribution of rare events including neonatal deaths [39]. Equation 1 presents the general mathematical form of Poisson multivariate regression model which was used in this study to undertake bivariate and multivariate regressions analysis.

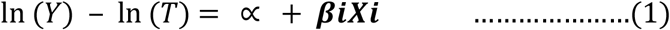

Where:-

Y is neonatal age segment total death counts
T is neonatal age segment total exposure days
*α* is intercept term
***X_i_*** is a vector of regression variables
***β_i_*** is a vector of regression coefficients

The dependent variable was specified as the natural logarithm of total number of deaths in age segment. The off-set variable was specified as the natural logarithm of total days of exposures in age segment. The vector of regression variables were the specified explanatory variables. The Generalized Linear Models (GLIM) subroutine in SPSS computer software package was used to fit bivariate and multivariate regression models. It provided parameter estimates for covariate odds ratios in the regression equations. Since odds ratio of a variable reference category is one (1.000), it then follows that ratios greater than 1 indicate increased likelihood of mortality and those ratios less than 1 indicate reduced likelihood of mortality relative to the reference category. Only odds ratios with at least 95% statistical significance level were considered to have significant effects in this analysis.

## Results

### Neonatal age segments mortality differentials in Kenya’s malaria prone areas by study ANC services

Out of the total 20794 births analysed in this study, there are 440 neonatal deaths which are distributed as follows: 33% in age segment less than 1 day; 54% in age segment 1 to 7 days; and, 13% in age segment 8 days to less than one month. The computed life table probabilities of dying (per 1,000 live births) during neonatal age segments are: 7 by first day; 18 by first week; and, 21 before attaining age one month. Table 1 reports estimated neonatal period mortality rates, with 95% CI, for the study ANC services variables in Kenya’s malaria areas.

**Table 1.**
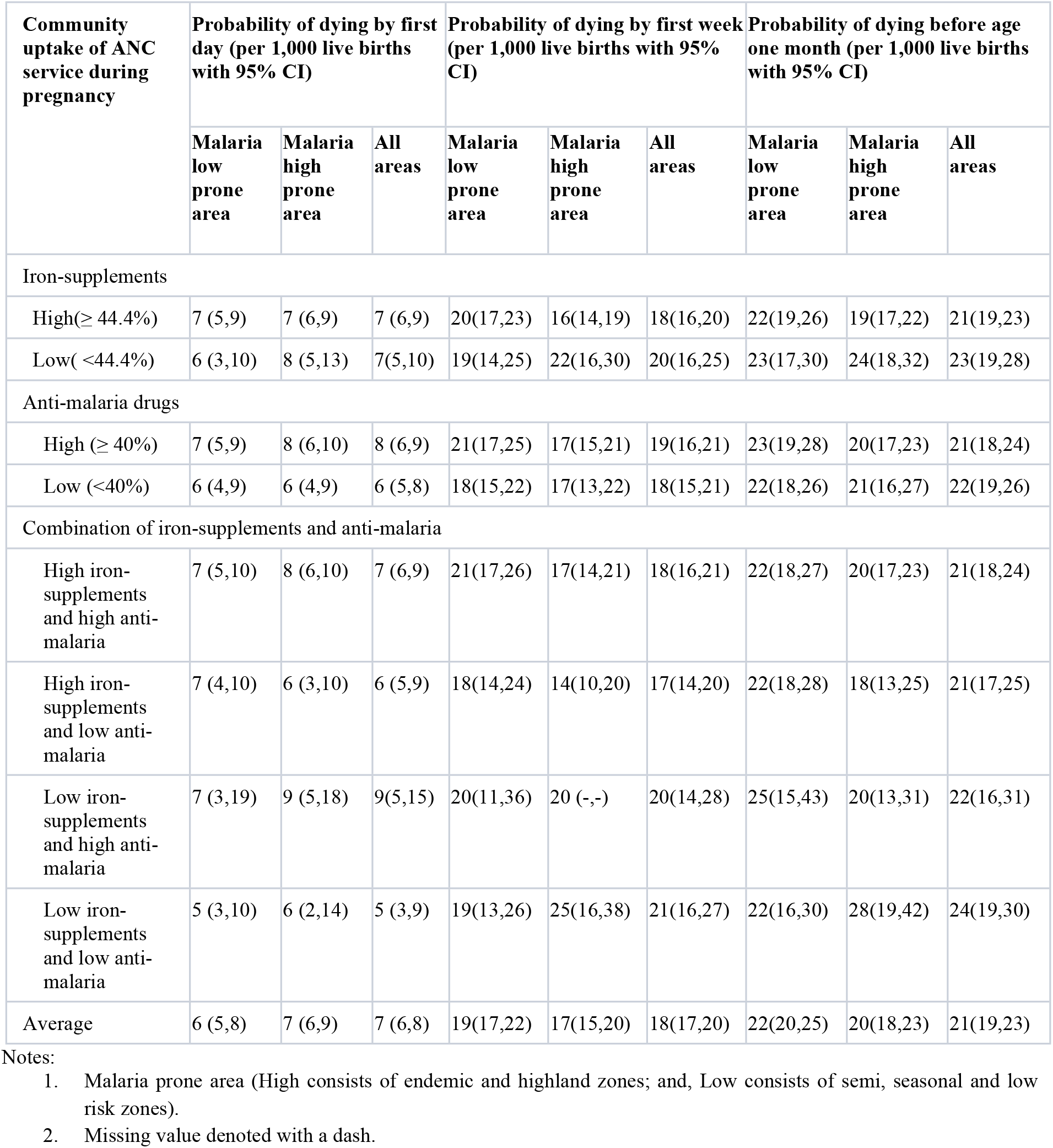
Neonatal age segments mortality rates for study ANC services variables in Kenya’s malaria areas.

The 95% confidence intervals for the mortality estimates indicate that there are no significant variations in age segment mortality rates for high and low malaria prone areas in Kenya. However, during the first day of life, high malaria prone areas have slightly higher mortality rates compared to low prone areas. On community uptake of iron-supplements during pregnancy, the results show that there are no significant variations in neonatal period age segments mortality rates in malaria areas. However, low community uptake of iron-supplements has higher mortality rates in all neonatal period age segments in high malaria prone areas relative to low malaria areas.

The results also show that there are no significant variations in neonatal period age segments mortality by community uptake of anti-malaria drugs during pregnancy in Kenya’s malaria areas. However, low community uptake of anti-malaria drugs has slightly higher mortality rates before attainment of age one month in high malaria prone areas (21 per 1,000 live births) compared with low malaria areas (20 per 1,000 live births).

Analysis results for various combinations of community uptake of iron-supplements and anti-malaria drugs during pregnancy show that there are no significant variations in neonatal period age segments mortality by combination of the uptake in Kenya’s malaria areas. However, the combination of low iron-supplements and low anti-malaria drugs compared to that of high iron-supplement and high anti-malaria drugs has much higher mortality rates during later two neonatal period age segments (first week and less than one month) in high malaria prone areas relative to their counterparts in low malaria areas. The combination of low iron-supplements and low anti-malaria drugs has probabilities of dying per 1,000 live births by ages one week and less than one month estimated at 25 and 28, respectively. The comparable mortality rates for combination of high iron-supplements and high anti-malaria drugs are 17 and 20, respectively.

### Effects of the study ANC services on risk of death during day of birth

Table 2 presents odds ratios with 95% CI and p-values for community uptake of iron-supplements and anti-malaria drugs during pregnancy variables on risk of death during the day of birth in Kenya’s malaria areas. The odds ratios were obtained from fitted three multivariate Poisson regression models (Model 1 for low malaria prone, Model 2 for high malaria prone and Model 3 for all areas). The table also provides odds ratios and p-values for interactive/combination variables involving uptake of the two study ANC services in Models 1 and 2. In addition, Table 2 presents odds ratios and p-values for three socio-demographic and birth characteristics variables (HIV positive pregnant women per 10,000 population; maternal age at child’s birth in years; and, birth type) and one variable associated with antenatal, delivery and post-delivery care services (proportion in community with early breastfeeding initiation period of less than 24 hours) that were used as adjustment variables in the three regression models fitted. Malaria prone area was only used as adjustment variable in Model 3.

**Table 2.** Odds ratios for mortality during day of birth for study ANC services and adjustment variables in Kenya’s malaria areas.

| Variable | Model 1<br>(Malaria low prone area) |  | Model 2<br>(Malaria high prone area) |  | Model 3 (All areas) |  |
| --- | --- | --- | --- | --- | --- | --- |
|  | Odds ratio ( 95% CI) | p-value | Odds ratio ( 95% CI) | p-value | Odds ratio ( 95% CI) | p-value |
| Community uptake of ANC service during pregnancy |  |  |  |  |  |  |
| Iron-supplements |  |  |  |  |  |  |
| High ($\geq 44.4\%$) | 0.639 (0.278-1.469) | 0.292 | 1.072 (0.379-3.034) | 0.896 | 0.934 (0.501- 1.742) | 0.830 |
| Low ($<44.4\%$) | 1.000 | | 1.000 | | 1.000 | |
| Anti-malaria drugs |  |  |  |  |  |  |
| High ($\geq 40\%$) | 1.962 (0.594-6.477) | 0.269 | 1.504 (0.437-5.182) | 0.518 | 1.842 (0.837-4.057) | 0.129 |
| Low ($<40\%$) | 1.000 | | 1.000 | | 1.000 | |
| Combination of iron-supplements and anti-malaria |  |  |  |  |  |  |
| High iron-supplements and high anti-malaria | 0.567 (0.144-2.235) | 0.417 | 0.523 (0.144-1.907) | 0.327 | 0.463 (0.196-1.093) | 0.079* |
| High iron-supplements and low anti-malaria | 1.000 |  | 1.000 |  | 1.000 |  |
| Low iron-supplements and high anti-malaria | 1.000 |  | 1.000 |  | 1.000 |  |
| Low iron-supplements and low anti-malaria | 1.000 |  | 1.000 |  | 1.000 |  |
| Adjustment variables |  |  |  |  |  |  |
| Maternal age at child's birth (in years) |  |  |  |  |  |  |
| 18-34 | 0.446 (0.247-0.804) | 0.007** | 0.574 (0.337-0.977) | 0.041** | 0.519 (0.356-0.757) | 0.001** |
| <18 and 35+ | 1.000 |  | 1.000 |  | 1.000 |  |
| Birth type |  |  |  |  |  |  |
| Single | 0.040 (0.018-0.085) | 0.000*** | 0.200 (0.095-0.421) | 0.000*** | 0.110 (0.066-0.186) | 0.000*** |
| Multiple | 1.000 |  | 1.000 |  | 1.000 |  |
| Breastfeeding initiation in community (proportion with < 24 hours) |  |  |  |  |  |  |
| High ($\geq 62.5\%$) | 0.175 (0.072-0.423) | 0.000*** | 0.189 (0.073-0.488) | 0.001** | 0.190 (0.102-0.354) | 0.000*** |
| Low ($<62.5\%$) | 1.000 | | 1.000 | | 1.000 | |
| HIV positive pregnant women (per 10,000 population) |  |  |  |  |  |  |
| Low ($<14.4$) | 0.592 (0.328-1.089) | 0.082* | 0.596 (0.285-1.245) | 0.169 | 0.629 (0.420-0.941) | 0.024** |
| High ($\geq 14.4$) | 1.000 | | 1.000 | | 1.000 | |
| Malaria area |  |  |  |  |  |  |
| Low prone |  |  |  |  | 1.187 (0.819-1.719) | 0.365 |
| High prone |  |  |  |  | 1.000 |  |

| Model parameters |  |  |  |  |  |  |
| --- | --- | --- | --- | --- | --- | --- |
| Intercept term | 3.416 (0.676-17.266) | 0.137 | 0.441 (0.059-2.867) | 0.370 | 0.752(0.244-2.318) | 0.620 |
| Likelihood ratio $\chi^2$ | 61.264 | | 25.477 | | 77.015 | |
| Degrees of freedom | 7 |  | 7 |  | 8 |  |
| p-value | 0.000*** |  | 0.000*** |  | 0.000*** |  |

Notes:
1. Malaria prone area (High consists of endemic and highland zones; and, Low consists of semi, seasonal and low risk zones) 2. \*\*\*p < 0.001, \*\*p < 0.05 and \* p < 0.1

The results show that only combined high iron-supplements and high anti-malaria drugs community uptake during pregnancy has significant reduction effect on risk of neonatal death in the first day of life but at 90% significant level. The results also show that all the four mortality risk factors which were adjusted for in the fitted regression models are significant mortality risk factors during the day of birth in both low and high malaria areas in Kenya.

### Effects of the study ANC services on risk of death during the period 1 to 7 days

Table 3 provides odds ratios with 95% CI and p-values for community uptake of iron-supplements and anti-malaria drugs during pregnancy variables on risk of child death during the period 1 to 7 days in Kenya’s malaria areas. The strategy used to undertake regression analysis for the first neonatal age segment (day of birth) was replicated in this second age segment. However, the only difference was on the adjustment variables used. Five socio-demographic and birth characteristics variables (HIV positive pregnant women per 10,000 population; preceding birth interval length in months; birth type; elevated birth order risk; and, maternal education) were used as adjustment variables in the three regression models fitted. Two variables associated with antenatal, delivery and post-delivery care services (proportion in community with required tetanus injections during pregnancy; and, mode of delivery) were also included as control variables in the three regression models.

**Table 3.** Odds ratios for mortality during the period 1 to 7 days for study ANC services and adjustment variables in Kenya’s malaria areas.

| Variable | Model 1<br>(Malaria low prone area) |  | Model 2<br>(Malaria high prone area) |  | Model 3 (All areas) |  |
| --- | --- | --- | --- | --- | --- | --- |
|  | Odds ratio (95% CI) | p-value | Odds ratio (95% CI) | p-value | Odds ratio (95% CI) | p-value |
| Community uptake of ANC service during pregnancy |  |  |  |  |  |  |
| Iron-supplements |  |  |  |  |  |  |
| High ($\geq 44.4\%$) | 0.730 (0.424-1.257) | 0.256 | 0.445 (0.216-0.919) | 0.028** | 0.587 (0.384-0.898) | 0.014** |
| Low ($<44.4\%$) | 1.000 | | 1.000 | | 1.000 | |
| Anti-malaria drugs |  |  |  |  |  |  |
| High ($\geq 40\%$) | 1.848 (0.772-4.426) | 0.168 | 0.669 (0.248-1.801) | 0.426 | 1.360 (0.723-2.558) | 0.340 |
| Low ($<40\%$) | 1.000 | | 1.000 | | 1.000 | |
| Combination of iron-supplements and anti-malaria |  |  |  |  |  |  |
| High iron-supplements and high anti-malaria | 0.585 (0.223-1.537) | 0.277 | 0.865 (0.298-2.514) | 0.790 | 0.680(0.342-1.353) | 0.272 |
| High iron-supplements and low anti-malaria | 1.000 |  | 1.000 |  | 1.000 |  |
| Low iron-supplements and high anti-malaria | 1.000 |  | 1.000 |  | 1.000 |  |
| Low iron-supplements and low anti-malaria | 1.000 |  | 1.000 |  | 1.000 |  |
| Adjustment variables |  |  |  |  |  |  |
| Mode of delivery |  |  |  |  |  |  |
| Not caesarean section | 0.290 (0.156-0.540) | 0.000*** | 0.213 (0.084-0.536) | 0.001** | 0.304 (0.186-0.497) | 0.000*** |
| Caesarean section | 1.000 |  | 1.000 |  | 1.000 |  |
| Preceding birth interval length (in months) |  |  |  |  |  |  |
| 12+ | 0.044 (0.018-0.113) | 0.000*** | 0.065 (0.020-0.211) | 0.000*** | 0.053 (0.026-0.111) | 0.000*** |
| $<12$ | 1.000 | | 1.000 | | 1.000 | |
| Elevated birth order risk |  |  |  |  |  |  |
| Low (2-3 & mother age 18-34) | 1.098 (0.744-1.621) | 0.636 | 1.112 (0.719-1.720) | 0.633 | 1.105 (0.827-1.476) | 0.499 |
| High (1,4+ & mother age $<18,35+$) | 1.000 | | 1.000 | | 1.000 | |
| Birth type |  |  |  |  |  |  |
| Single | 0.115 (0.059-0.223) | 0.000*** | 0.060 (0.035-0.102) | 0.000*** | 0.079 (0.053-0.117) | 0.000*** |
| Multiple | 1.000 |  | 1.000 |  | 1.000 |  |
| Mother's education level |  |  |  |  |  |  |
| Secondary and above | 0.991 (0.581-1.691) | 0.975 | 1.135 (0.678-1.898) | 0.630 | 1.122 (0.779-1.617) | 0.537 |
| None/primary | 1.000 |  | 1.000 |  | 1.000 |  |

| Child tetanus protection (community uptake of required injections during pregnancy) |  |  |  |  |  |  |
| --- | --- | --- | --- | --- | --- | --- |
| High ($\geq 55.6\%$) | 1.020 (0.681-1.527) | 0.925 | 1.170 (0.743-1.843) | 0.497 | 1.063 (0.787-1.436) | 0.688 |
| Low ($< 55.6\%$) | 1.000 | | 1.000 | | 1.000 | |
| HIV positive pregnant women (per 10,000 population) |  |  |  |  |  |  |
| Low ($< 14.4$) | 0.808 (0.532-1.227) | 0.317 | 0.789 (0.483-1.344) | 0.384 | 0.943 (0.690-1.290) | 0.716 |
| High ($\geq 14.4$) | 1.000 | | 1.000 | | 1.000 | |
| Malaria area |  |  |  |  |  |  |
| Low prone |  |  |  |  | 1.218 (0.908-1.634) | 0.188 |
| High prone |  |  |  |  | 1.000 |  |
| Model parameters |  |  |  |  |  |  |
| Intercept term | 1.785 (.476-6.696) | 0.390 | 4.663(.789-27.560) | 0.089* | 1.767(0.629-4.964) | 0.280 |
| Log Likelihood $\chi^2$ | 78.824 | | 103.163 | | 172.502 | |
| Degrees of freedom | 10 |  | 10 |  | 11 |  |
| p-value | 0.000*** |  | 0.000*** |  | 0.000*** |  |

Notes:
1. Malaria prone area (High consists of endemic and highland zones; and, Low consists of semi, seasonal and low risk zones) 2. \*\*\*p < 0.001, \*\*p < 0.05 and \* p < 0.1

The results show that high community uptake of iron-supplements during pregnancy has significant reduction effect on risk of child death during the neonatal period 1 to 7 days in Model 3 (for combined low and high malaria prone areas). The results also indicate that only two socio-demographic and birth characteristics variables (preceding birth interval length in months; and, birth type) and only one variable associated with antenatal, delivery and post-delivery care services (mode of delivery) are significant control variables in all the three fitted regression models.

### Effects of the study ANC services on risk of death during the period 8 days to less than a month

Table 4 presents odds ratios with 95% CI and p-values for community uptake of iron-supplements and anti-malaria drugs during pregnancy variables on risk of child death during the period 8 days to less than a month in Kenya’s malaria prone areas. The regression analysis strategy applied in this third neonatal age segment was similar to the one used in the first two age segments. However, two socio-demographic and birth characteristics variables (HIV positive pregnant women per 10,000 population; and, birth type) were used as adjustment variables in the three regression models fitted. In addition, one variable associated with antenatal, delivery and post-delivery care services (proportion of children sick with diarrhoea in community who received appropriate care) was included as a control variable in the three regression models fitted.

**Table 4.**
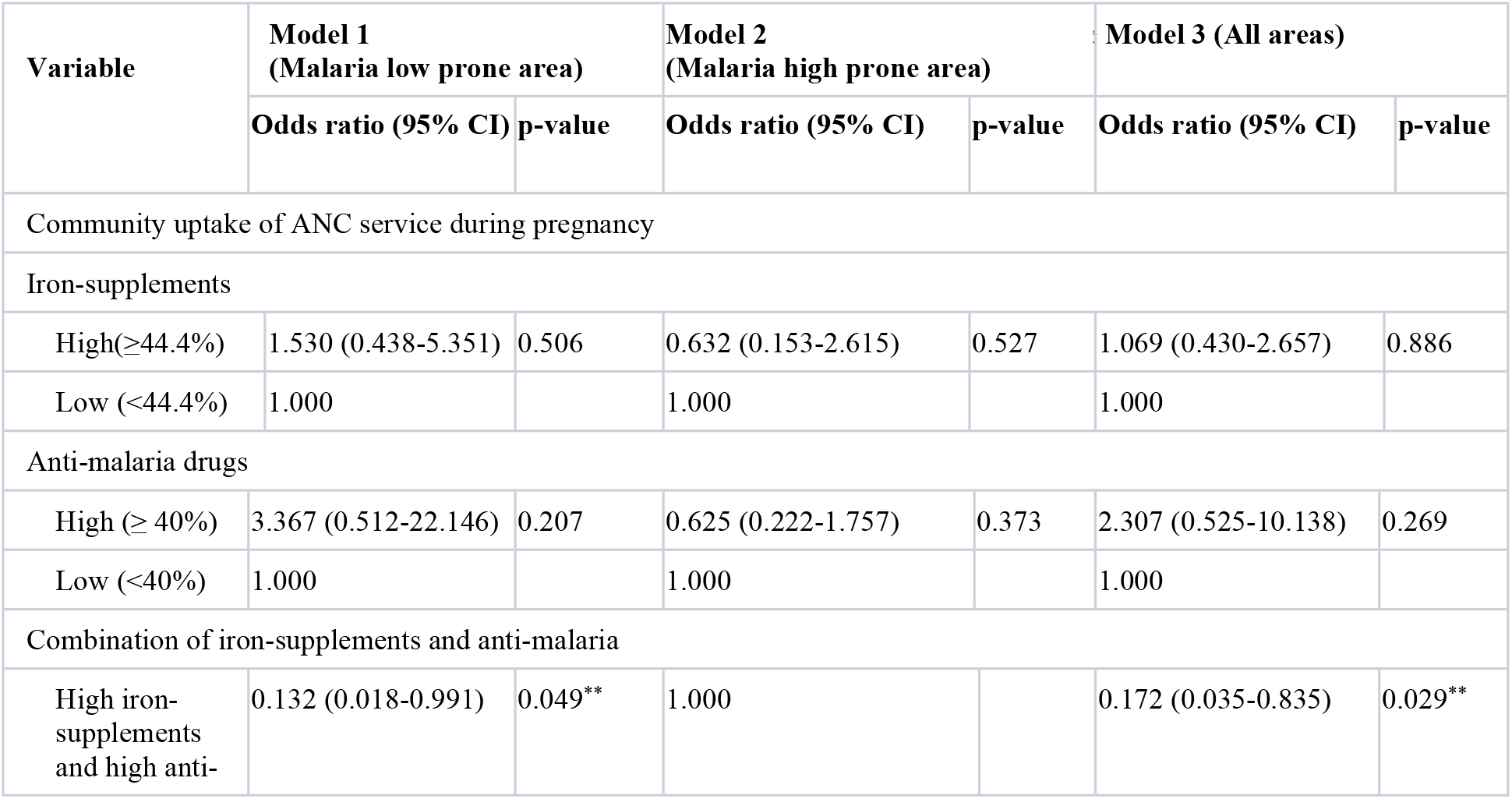
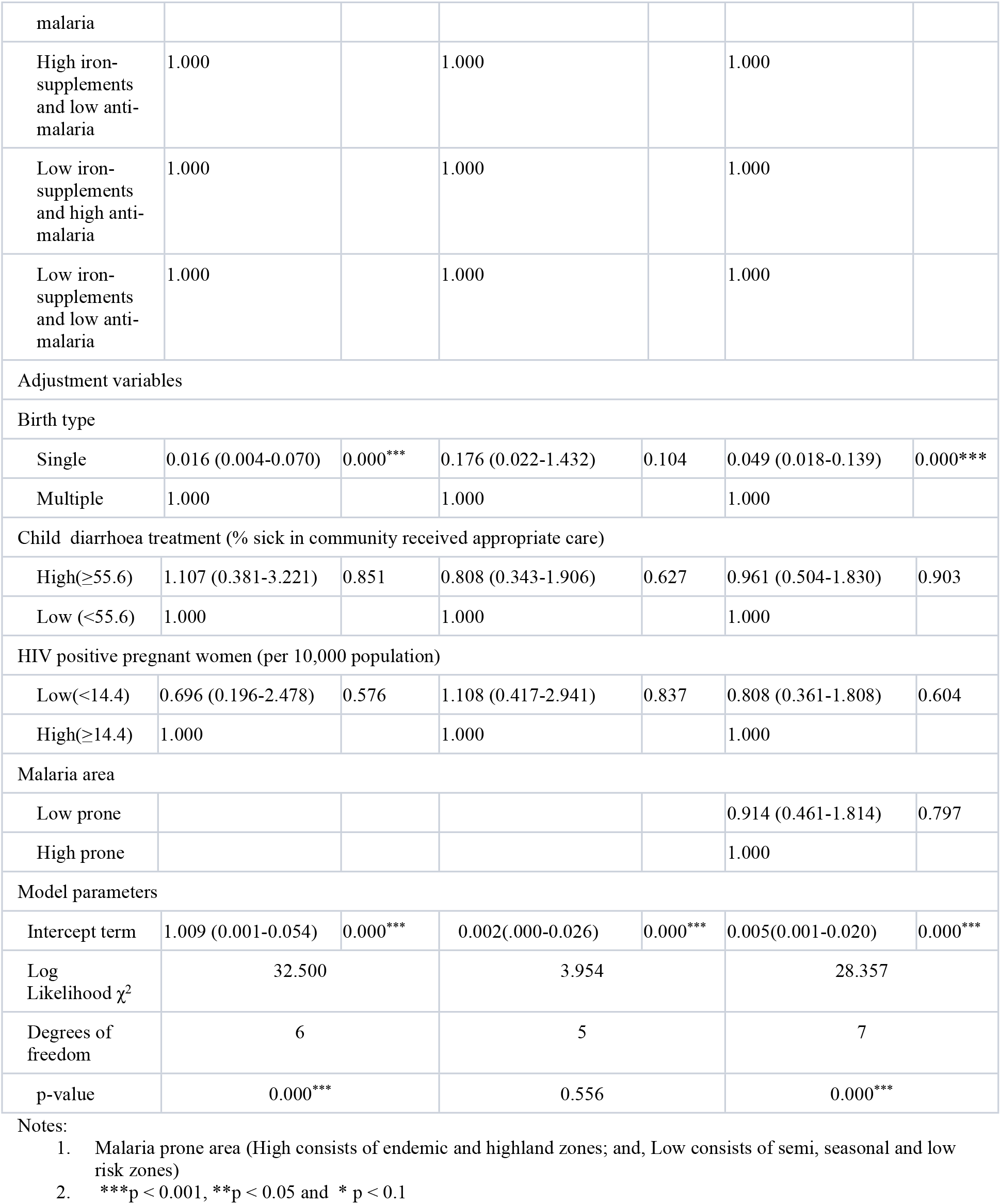
Odds ratios for mortality during the period 8 days to less than a month for study ANC services and adjustment variables in Kenya’s malaria areas.

The results show that the combination of community uptake of high iron-supplements with high iron-supplements during pregnancy has significant reduction effect on risk of death during the period 8 days to less than a month in Model 1 (malaria low prone area) and Model 3 (all areas). The results also indicate that only one socio-demographic and birth characteristics control variable (birth type) is significant in Models 1 and 3. In addition, the only one control variable associated with antenatal, delivery and post-delivery care services (proportion of children sick with diarrhoea in community who received appropriate care), included in this age segment analysis, is not significant.

## Discussion

This study investigates variations in mortality in three neonatal period age segments in epidemiological malaria areas in Kenya by community uptake of two ANC services during pregnancy with the aim of determining the effects of uptake of these services on age segment mortality. The results depict insignificant variations in age segment mortality rates by the two study ANC services, but a mixed picture of the beneficial effects of these services on mortality risks in the neonatal age segments.

The study results on age pattern of mortality during neonatal period in Kenya are consistent with the global, south Asia and sub-Saharan Africa regions patterns [2,8,30]. The analysis show that in both high and low malaria prone areas, the force of mortality is greatest in the second age segment (1 to 7 days) followed by the first segment (day of birth) and lowest in the last age segment (8 days to less than one month). Based on the mortality rates obtained, the contribution of early neonatal mortality (0 to 7 days) to Kenya’s neonatal mortality rate is about 75%. Its contribution to neonatal mortality rates in low malaria prone areas and high malaria areas are 80% and 100%, respectively.

Although the results on mortality rates variations in neonatal age segments are statistically insignificant, the depicted patterns are consistent with the study expectations. Low uptake of the study ANC services treated separately and in combination is associated with higher mortality rates in all age segments in high malaria prone areas in Kenya. The finding that uptake of high iron-supplements and high anti-malaria drugs during pregnancy reduces significantly mortality risks in the third age segment (8 days to less than one month) relative to low iron-supplements and low anti-malaria uptake is generally consistent with earlier findings for south Asia and sub-Saharan Africa regions [14,16,24].

The unexpected finding, although statistically insignificant, is that high community uptake of anti-malaria drugs during pregnancy is associated with high neonatal period age segments mortality. This stands in contrast to previous studies, especially with the south Asia study, which indicated that intermittent preventive treatment of malaria in pregnancy is associated with about 31% reduction in neonatal mortality [10]. A possible explanation is that in Kenya, high malaria areas are also high childhood mortality areas and use of anti-malaria drugs for prevention and treatment in the general population is common. In addition, provision of anti-malaria drugs during pregnancy is not restricted to ANC services but can be obtained readily without qualified health personnel’s prescription in non-health service outlets including local shops/kiosks. The reported uptake of anti-malaria drugs during pregnancy in 2014 KDHS may have captured uptake of anti-malaria drugs by mothers beyond pregnancy durations.

This study also identifies and provides effects of the mortality risk factors which were controlled for in analysing death risks in neonatal age segments in Kenya’s malaria areas. The findings show that the number of significant factors associated with broad group of socio-demographic and birth characteristics as well as those associated with antenatal, delivery and post-delivery care services, declined after early neonatal (0 to 7 days). This suggests that in Kenya, further reduction in mortality risk in late neonatal (8 days to less than one month) may possibly depend largely on care for small and sick neonates. This possibility is not tested in this analysis but is deduced from the WHO model which attributes neonatal deaths to readily preventable and treatable causes with cost-effective interventions covering the antenatal period, time around birth and first week of life and care for small and sick neonates [2,7,8].

Due to poor quality of data collected in 2014 KDHS on care for small and sick neonates, their effects on late neonatal mortality could not be examined in this study. The 2014 KDHS gathered information on immunization, health and nutrition status of all under five year old children born to women who were interviewed [3]. In this study, measurements for variables on care and sick neonates were assumed not to differ substantially from those for the under five year old children. Only one variable on reported care given to sick children with diarrhoea qualified for inclusion as control in the final fitted regression models. The other three variables (care given to child with fever/cough, child faecal disposal and timely postnatal care for non-facility delivery births), which were initially considered for inclusion in this study, were found to be insignificant at bivariate analysis stage.

## Limitations

This study has limitations which may have affected the accuracy of the computed neonatal age segments mortality. First, is the missing information on age in completed days for all neonates found alive on interview date and whose individual age in days were assumed to be 15 in this analysis. Second, is the missing information on maternal uptake of the study ANC services for any child in the dataset who was not a last-born or reported dead on interview date. This necessitated use of weighted cluster level proportions. Third, this study is based on cross-sectional data. Cross-sectional data are subject to omissions and misreporting of information. This analysis used information on individual children born to women during the period 0 to 59 months before the interview, excluding interview month. This meant that some of the reported dates of events, especially on uptake of study ANC services during last pregnancy, could have been misplaced taking into consideration the possible maximum 59 months duration prior to interview date.

## Conclusions

This study establishes that variations in neonatal age segments mortality rates by study ANC services in malaria epidemiological areas in Kenya, using cross-sectional 2014 KDHS data, are not statistically significant. It shows that early neonatal contributes about 80% and above of neonatal mortality in all Kenya’s malaria zones. The analysis provides effects of iron-supplements and ant-malaria drugs uptake during pregnancy as well as other mortality risk factors used as control variables on neonatal age segments mortality. The study also establishes that combination of high uptake of iron-supplements and high anti-malaria drugs during pregnancy compared to combination of low iron-supplements and low anti-malaria uptake, reduce significantly risk of death in late neonatal period of 8 days to less than one month in low and all malaria areas in Kenya. The study findings have implications for neonatal survival programmes implementation and future KDHS data collection in Kenya. Efforts to attain near universal uptake (99%) for the study ANC services should be intensified given that their 2014 cut-off points for high community uptake during pregnancy are both below 50% (44% for iron-supplements and 40% for anti-malaria drugs). This study recommends that efforts be made to improve quality of data on care for small and sick neonates collected in future DHS type cross-sectional surveys in Kenya.

## Acknowledgements

The author would like to thank the Kenya National Bureau of Statistics (KNBS) for availing the 2014 KDHS data and the sampled districts codes which facilitated computation of community variables and categorization of the districts into malaria areas based on the 2015 Kenya MALARIA Indicator Survey Report Appendix A. Comments and suggestions from reviewers of this manuscript are also acknowledged in advance.

